# Identifying hetero-protein complexes in the nuclear envelope

**DOI:** 10.1101/709196

**Authors:** J. Hennen, K.H. Hur, J. Kohler, S.R. Karuka, I. Angert, G.W.G. Luxton, J.D. Mueller

## Abstract

The nucleus is delineated by the nuclear envelope (NE), which is a double membrane barrier composed of the inner and outer nuclear membranes as well as a ~40 nm wide lumen. In addition to its barrier function, the NE acts as a critical signaling node for a variety of cellular processes which are mediated by protein complexes within this subcellular compartment. While fluorescence fluctuation spectroscopy (FFS) is a powerful tool for characterizing protein complexes in living cells, it was recently demonstrated that conventional FFS methods are not suitable for applications in the NE because of the presence of slow nuclear membrane undulations. We previously addressed this challenge by developing time-shifted mean-segmented Q (tsMSQ) analysis and applied it to successfully characterize protein homo-oligomerization in the NE. However, many NE complexes, such as the linker of the nucleoskeleton and cytoskeleton (LINC) complex, are formed by heterotypic interactions, which single-color tsMSQ is unable to characterize. Here, we describe the development of dual-color (DC) tsMSQ to analyze NE hetero-protein complexes built from proteins that carry two spectrally distinct fluorescent labels. Experiments performed on model systems demonstrate that DC tsMSQ properly identifies hetero-protein complexes and their stoichiometry in the NE by accounting for spectral crosstalk and local volume fluctuations. Finally, we applied DC tsMSQ to study the assembly of the LINC complex, a hetero-protein complex composed of Klarsicht/ANC-1/SYNE homology (KASH) and Sad1/UNC-84 (SUN) proteins, in the NE of living cells. Using DC tsMSQ, we demonstrate the ability of the SUN protein SUN2 and the KASH protein nesprin-2 to form a hetero-complex *in vivo*. Our results are consistent with previously published *in vitro* studies and demonstrate the utility of the DC tsMSQ technique for characterizing NE hetero-protein complexes.

**Statement of Significance:** Protein complexes found within the nuclear envelope (NE) play a vital role in regulating cellular functions ranging from gene expression to cellular movement. However, the assembly state of these complexes within their native environment remains poorly understood, which is compounded by a general lack of fluorescence techniques suitable for quantifying the oligomeric state of NE protein complexes. This study aims at addressing this issue by introducing dual-color time-shifted mean-segmented Q analysis as a fluorescence fluctuation method specifically designed to identify the average oligomeric state of hetero-protein complexes within the NE of living cells.

## Introduction

Fluorescence fluctuation spectroscopy (FFS) refers to a collection of related biophysical techniques that exploit the stochastic intensity of fluorescently labeled biomolecules passing through a small observation volume created by confocal or two-photon microscopy [1]. The primary parameters accessible by FFS are the concentration, mobility, and oligomeric state of the labeled biomolecule [1]. While the original analysis of FFS results was based on the autocorrelation function (ACF), many other methods have been introduced over the years, each with its own strengths and weaknesses [2–5]. An important advance in FFS was the introduction of dual-color (DC) FFS for identifying interactions between two species of biomolecules labeled with spectrally distinct fluorophores [6]. In DC FFS, the emission of the fluorophores is separated by color into two detection channels. Heterotypic interactions between the two species lead to synchronized temporal fluctuations in both channels, which are recognized by the cross-correlation function (CCF) of the two detected signals. In addition to CCF, other analysis techniques have been introduced for quantifying hetero-species interactions from FFS experiments [7,8].

Because FFS is an equilibrium technique that passively observes fluctuations, it is well suited for applications in live cells [9]. Cellular proteins can be conveniently labeled by genetic tagging with one of the many available fluorescent proteins [10]. Brightness, which characterizes the intrinsic fluorescence intensity of a molecule, is an important FFS parameter because it contains information about the stoichiometry of fluorescently-tagged protein complexes [11]. For example, the brightness of monomeric proteins tagged with EGFP will increase upon their association into homo-oligomers, as each protein complex contains several fluorescent labels. This concept has been generalized to include differently colored fluorophores to characterize hetero-protein complexes [7]. While FFS brightness analysis has been successfully used to quantify protein-protein interactions within the cytoplasm, nucleoplasm, and at the plasma membrane [9,12,13], its extension to the nuclear envelope (NE) has proven challenging [14].

The NE consists of an inner and outer nuclear membrane (INM and ONM, respectively) separated by a ~40 nm thick fluid layer, known as the lumen or perinuclear space. Although the NE has been identified as a critical signaling node of the cell [15], a mechanistic understanding of how these hetero-protein complexes assemble remains limited due to the lack of quantitative biophysical techniques suitable for use in the NE. To address this challenge, we explored the use of single-color (SC) FFS for characterizing homo-protein association in the NE of living cells [14,16]. We found that slow undulations of the nuclear membranes give rise to local volume fluctuations that are not properly accounted for by conventional FFS methods [14]. We overcame this obstacle by introducing mean-segmented Q (MSQ) analysis as well as time-shifted MSQ (tsMSQ), a significantly improved version of MSQ [14,17,18]. We demonstrated that these techniques successfully identify the mobility and homo-oligomeric state of NE proteins [14].

This study extends FFS analysis of hetero-protein complexes to the NE by introducing DC tsMSQ, a generalized form of regular tsMSQ. The DC tsMSQ framework includes hetero-species partitioning (HSP) [7] and effectively eliminates complications due to spectral crosstalk in the characterization of heterotypic protein interactions as verified by control experiments. Our study demonstrates that local volume fluctuations of the NE are a significant challenge for conventional DC FFS analysis, which prompted the development of DC tsMSQ. We first verified the foundation of DC tsMSQ using pairs of interacting and non-interacting proteins measured in the cytoplasm and in the NE. In addition, control experiments using both luminal and nuclear membrane-associated proteins were conducted to illustrate the influence of the nuclear membrane undulations on DC tsMSQ analysis.

To demonstrate the power of DC tsMSQ, we applied it towards studying the assembly of the linker of nucleoskeleton and cytoskeleton (LINC) complex in the NE of living cells. This NE-spanning molecular bridge mediates mechanical force transmission into the nucleoplasm and is required for several fundamental cellular processes including cell division, DNA damage repair, meiotic chromosome pairing, mechano-regulation of gene expression, and nuclear positioning [19]. The LINC complex is formed by a direct transluminal heterotypic interaction between the cytoskeletal-binding ONM *Klarsicht*/*ANC-1*/*SYNE* homology (KASH) proteins and the nuclear lamina-binding INM *Sad1*/*UNC-84* (SUN) proteins [20]. Previous *in vitro* biochemical and structural studies revealed that the luminal domain of SUN2 homo-trimerizes and that the luminal domain of nesprin-2 binds in the grooves formed at the interface of two adjacent interacting SUN2 monomers [21,22]. Thus, the homo-trimerization of the SUN2 luminal domain enables the recruitment of 3 nesprin-2 luminal domains resulting in the assembly of a SUN2-nesprin-2 hetero-hexamer. We recently succeeded in directly measuring the homo-trimerization of the EGFP-tagged luminal domain of SUN2 in the NE of living cells by SC FFS [16]. Here we utilize DC tsMSQ to directly observe the heterotypic interactions formed between the EGFP-tagged SUN2 luminal domain and an mCherry-tagged construct that encodes the last three spectrin-like repeats (SRs), the transmembrane domain, and the luminal KASH peptide of nesprin-2 in the NE. This work establishes the theoretical and practical framework necessary for future quantitative studies of LINC complex assembly within the NE of living cells as well as the characterization of additional heterotypic interactions between proteins within this relatively unexplored subcellular environment.

## Material and Methods

### Experimental Setup

Brightness measurements were performed as previously described [7,23]. Briefly, FFS data were acquired on a custom built two-photon microscope with a 63x C-Apochromat water-immersion objective with numerical aperture (NA) = 1.2 (Zeiss, Oberkochen, Germany) using an excitation wavelength of 1000 nm and an average power after the objective in the range of 0.3 to 0.4 mW. A dichroic mirror centered at 580 nm (FF580-FDi01; Semrock, Rochester, NY) was used to split the emission path into two channels. An additional 84 nm wide bandpass filter centered at 510 nm (FF01-510/84; Semrock) was placed before the green channel to remove any reflected fluorescence from mCherry. Photon counts were detected using avalanche photodiodes (SPCM-AQ-141 APD; Perkin-Elmer, Dumberry, Quebec, Canada), recorded with a Flex04-12D card (correlator.com, Bridgewater, NJ) sampled at 20 kHz and analyzed using programs written in IDL 8.7 (Research Systems, Boulder, CO). Z-scans were performed using an arbitrary waveform generator (model No. 33522A; Agilent Technologies, Santa Clara, CA) to move a PZ2000 piezo stage (ASI, Eugene, OR) axially. The waveform created by the generator was a linear ramp function with a peak-to-peak amplitude of 1.6 V, corresponding to 24.1 μm of axial travel, and a period of 10 s for a speed of 4.82 μm/s.

### Measurement Procedure

EGFP (G) calibration measurements were performed in the cytoplasm of EGFP-expressing U2OS cells as previously described [12,23] to obtain its brightness in both the green and red channels (*λ*_*g,G*_ and *λ*_*r,G*_, respectively). Additional calibration measurements were performed in the cytoplasm of U2OS cells expressing EGFP-RARLBD-mCherry [7] to obtain the brightness *λ*_*g,Ch*_ of mCherry (Ch) in the red channel, which accounts for the previously described two-state brightness of this fluorescent protein [7]. These calibration values were then converted to *Q* values for the NE using *Q* = *γ*_2_*λT*_*S*_ where *γ*_2_ is the shape factor for a 2D Gaussian point spread function and *T*_*S*_ is the sampling time [12,24]. Measurements in the NE were performed as previously described [14,25] by first using epifluorescence to identify cells expressing the relevant constructs. FFS data were then acquired by taking z-scans through the nucleus which were analyzed as previously described [14,26]. Next, the two-photon beam was focused on the ventral NE followed by the dorsal NE and ~60 seconds of intensity fluctuation data were obtained at each location. These data were analyzed as described in the Theory section in order to obtain the normalized HSP brightness vector **b** = (*b*_*g*_, *b*_*r*_) [7,27]. The normalized brightness values were corrected for two-state brightness and fluorescence resonance energy transfer (FRET) as previously described [7,28]. The average number of molecules in the observation volume was determined by *N*_*g*_ = 〈*F*_*g*_〉/*λ*_*g,G*_ and *N*_*r*_ = (〈*F*_*r*_〉 − *f*_*ct*_ *〈*F*_*g*_〉)/*λ*_*r,Ch*_, where *f*_*ct*_ is the spectral crosstalk of EGFP given by *Q*_*r,G*_/*Q*_*g,G*_ [7].

### Sample Preparation

Experiments were conducted using transiently transfected U2OS cells (ATCC, Manassas, VA) maintained in DMEM with 10% fetal bovine serum (Hycolone Laboratories, Logan, UT). U2OS cells were subcultured into 24-well glass bottom plates (In Vitro Scientific, Sunnyvale, CA) prior to transfection. GenJet (SignaGen Laboratories, Rockville, MD) was used to transiently transfect cells 12-24 hours prior to measurement, according to the instructions of the manufacturer. The growth medium was replaced with Dulbecco’s phosphate-buffered saline containing calcium and magnesium (Biowhittaker, Walkerville, MD) immediately before measuring.

### Reagents

Restriction enzymes (REs) were either purchased from New England Biolabs (NEB, Ipswich, MA) or Promega (Madison, WI). Calf Intestinal Phosphatase, Phusion DNA polymerase, T4 DNA ligase, and T4 polynucleotide kinase were also purchased from NEB. All other chemicals were purchased from Sigma-Aldrich (St. Louis, MO) unless otherwise specified. The Wizard SV Gel and PCR Clean-Up System were purchased from Promega while the GeneJet Plasmid Midiprep Kit was purchased from ThermoFisher Scientific (Waltham, MA).

### DNA Constructs

The generation of SS-EGFP, SS-EGFP-torsinA^NTD-2xLeu^, SS-EGFP-SUN2^595-731^, and SS-EGFP-SUN2^261-731^ constructs were described previously [14,16]. The generation of SS-mCherry-KDEL, the SUN domain-linked SS-EGFP-SUN2^595-731^-mCherry (SS-EGFP-SL-mCherry), and mCherry-SR-KASH2 constructs are described in Text S6 of the Supporting Materials and Methods.

### Theory

#### Dual-channel tsMSQ_σρ_

The single-channel tsMSQ was generalized to dual-channel tsMSQ_σρ_ (Text S1 of the Supporting Materials and Methods) with the subscripts *σ* and *ρ* specifying the detection channel. A single diffusing species is described by

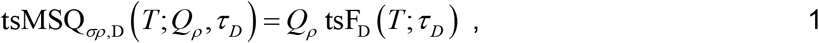

while the exponential correlation process caused by the local volume fluctuations of the NE is given by

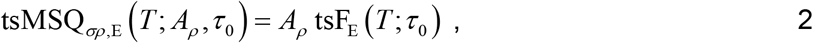

as derived in Text S2 of the Supporting Materials and Methods. A sample consisting of a single diffusing species in the lumen of the NE experiences local volume fluctuations and is described by the addition of Eqs. 1 and 2 [14],

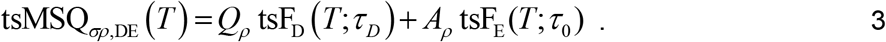

The time-shifted correlation functions tsF_D_ and tsF_E_ are defined in terms of second-order binning functions B_2,D_(*T*) and B_2,E_(*T*) [4,14,18,29]. Explicit formulas are found in Text S3 of the Supporting Materials and Methods. Since Eqs. 1 and 2 must reproduce the single channel case for *σ* = *ρ*, the amplitudes *Q*_*ρ*_ and *A*_*ρ*_ are given by *Q*_*ρ*_ = *γ*_2_*λ*_*ρ*_*T*_*s*_ and *A*_*ρ*_ = *c*^2^*λ*_*ρ*_*T*_*s*_*N*, respectively [14,18]. The brightness of the fluorescent molecule in the *ρ* channel is *λ*_*ρ*_, the number of molecules in the observation volume is *N*, and the shape factor of the observation volume is *γ*_2_. The factor *c* is determined by the fluctuations in the gap distance *h* separating the INM and ONM, 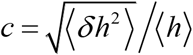 [14]. The diffusion time and the characteristic time of the volume fluctuations are given by *τ*_*D*_ and *τ*_0_, respectively.

The two detection channels used for DC FFS are labeled as green (g) and red (r). We define the DC tsMSQ function as

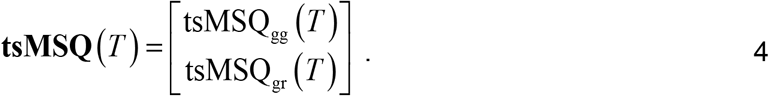

We refer to tsMSQ_gg_ as the autocorrelation tsMSQ of the green detection channel and tsMSQ_gr_ as the cross-correlation tsMSQ of the green and red channel. Thus, DC tsMSQ of a diffusing species is described by

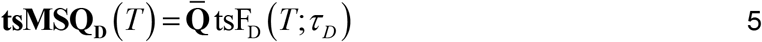

and in the presence of volume fluctuations the exponential correlation process

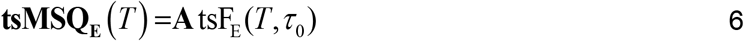

is added, resulting in

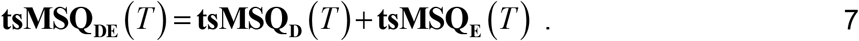

DC tsMSQ reduces to the same functional form as SC tsMSQ but with vector amplitudes 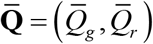 and **A** = (*A*_*g*_, *A*_*r*_). Finally, for a fit to a model with *S* diffusing species the **tsMSQ_D_** term is replaced by

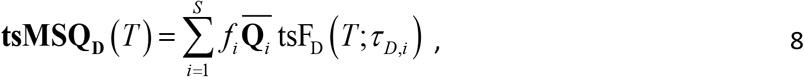

where 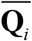 and *τ*_*D,i*_ are the are the Q-vector and diffusion time of the *i*^th^ species, respectively. The fractional intensity *f*_*i*_ of the *i*^th^ species is defined by the ratio of the green-channel intensity of *i*^th^ species to the total intensity 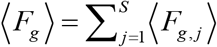 of the green channel, *f*_*i*_ = 〈*F*_*g,i*_〉/〈*F*_*g*_〉, as described in the Supporting Materials and Methods. The total Q-vector of the sample is defined by

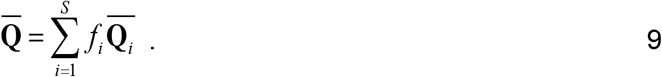

#### DC tsMSQ and HSP

We previously demonstrated that the complications in interpreting the results of FFS experiments in the presence of spectral crosstalk are avoided by HSP analysis [7]. HSP requires that there is no spectral leakage of the red-emitting mCherry into the green emission channel, which is accomplished by choosing appropriate filters [7]. Thus, the two fluorescent proteins EGFP (G) and mCherry (Ch) are characterized by their respective Q-vectors **Q**_*G*_ = (*Q*_*g,G*_, *Q*_*r,G*_) and **Q**_*ch*_ = (0,*Q*_*r,Ch*_). The Q-vector (Eq. 9) determined by DC tsMSQ analysis may be expressed as a linear combination of the Q-vectors of EGFP and mCherry,

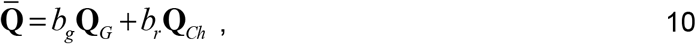

where the coefficients *b*_*g*_ and *b*_*r*_ represent the normalized brightness associated with the corresponding fluorescent proteins [7,30]. The tuple (*b*_*g*_, *b*_*r*_) characterizes the brightness vector of the hetero-species present in the sample. The hetero-species comprises all EGFP-labeled proteins and hetero-protein complexes carrying both EGFP and mCherry. Monomeric or oligomeric complexes that only contain the mCherry label are partitioned out by HSP. For example, a non-interacting monomeric EGFP-labeled species is described by a HSP brightness vector of (1, 0), while a hetero-dimer containing one EGFP and one mCherry label are described by a HSP brightness vector of (1, 1). A HSP brightness vector of (1, *y*) describes an EGFP-labeled protein that on average is associated with *y* mCherry-labeled proteins [7]. A graphical representation of these examples is provided in Fig. S1. The HSP brightness vector (*b*_*g*_, *b*_*r*_) is affected by fluorescent labels with dark states and multiple brightness states as well as by FRET, which bias the interpretation [7,28]. These effects were accounted and corrected for as previously described [7,28]. A derivation of HSP for DC tsMSQ is found in Text S5 of the Supporting Material and Methods.

## Results

Initial experiments were performed on a non-interacting pair of proteins located in the lumen of the NE (Fig. 1A). Specifically, we used SS-EGFP and SS-mCherry-KDEL, which have been found to be monomeric proteins within the NE by SC FFS (Fig. S2) [14]. EGFP and mCherry were chosen as labels because this pair has been characterized extensively by DC FFS and have a wide separation in their emission color [7,10]. The signal sequence (SS) of the luminal ATPase torsinA, which is cleaved after protein expression, ensures the presence of EGFP in the lumen of the NE [31]. The C-terminus of SS-mCherry was additionally fused to the endoplasmic reticulum retention signal KDEL to ensure its efficient targeting to the lumen. DC FFS data were collected at the NE of U2OS cells co-expressing SS-EGFP and SS-mCherry-KDEL. The ACFs of the fluorescence collected by the green and red detection channels were calculated (*G*_*gg*_ and *G*_*rr*_, respectively) as well as the cross-correlation (CCF) *G*_*gr*_ of both channels. We observed a positive CCF amplitude (Fig. 2), which was not unexpected for a non-interacting pair of fluorescent proteins, since spectral crosstalk leads to a positive CCF component [32]. This crosstalk-induced CCF function is predicted by [32]

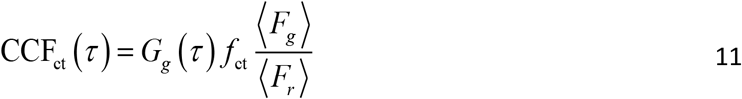

With 〈*F*_*g*_〉 and 〈*F*_*r*_〉 representing the mean fluorescence intensities of the green and red channel, respectively. The crosstalk in our setup is caused by the detection of the long wavelength emission of EGFP in the red detection channel [33], which is characterized by the crosstalk intensity fraction *f*_ct_. The observed CCF has to significantly exceed this baseline for a positive identification of heterotypic protein interactions [32]. The computed CCF_ct_(*τ*) (solid blue lines, Fig. 2) is not significantly different from the observed CCF for the data taken at low expression levels (Fig. 2A), but is significantly below the experimental CCF at medium (Fig. 2B) and high expression levels (Fig. 2C). This observation is counterintuitive, as it suggests the onset of interactions between EGFP and mCherry-KDEL at higher concentrations. We further noticed a significant change in the shape of the ACF of both channels as well as the CCF with increasing concentration (Fig. 2), which was unexpected because the diffusion process is independent of concentration. These changes in the ACF are specific to the NE environment [14] and are not observed in the cytoplasm, as confirmed by a control experiment performed in the cytoplasm of cells expressing EGFP and mCherry. Specifically, measurements performed in these cells showed a concentration-independent shape of the ACF as well as a strong overlap between the computed CCF_ct_(*τ*) and the CCF over a wide range of expression levels (Fig. S3), indicating the absence of significant interactions between EGFP and mCherry within the cytoplasm.

**Figure 1.**
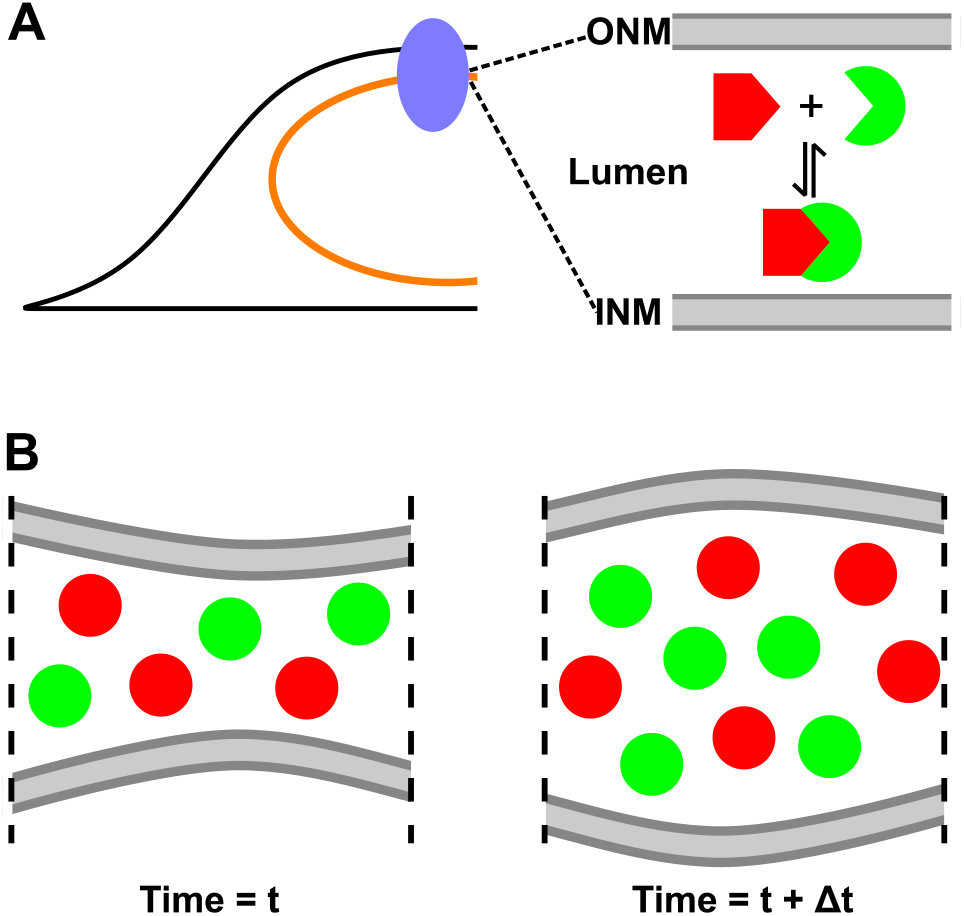
FFS at the NE. A) Illustration of a cell expressing potentially interacting green and red fluorescently labeled NE proteins with the two-photon excitation volume (blue oval) focused at its NE (orange circle), which consists of the INM and ONM separated by a ~40 nm thick lumen. B) Illustration of the time-dependent local volume fluctuations caused by nuclear membrane undulations, which give rise to coupled intensity variations of the non-interacting green and red fluorescently labeled proteins.

**Figure 2.**
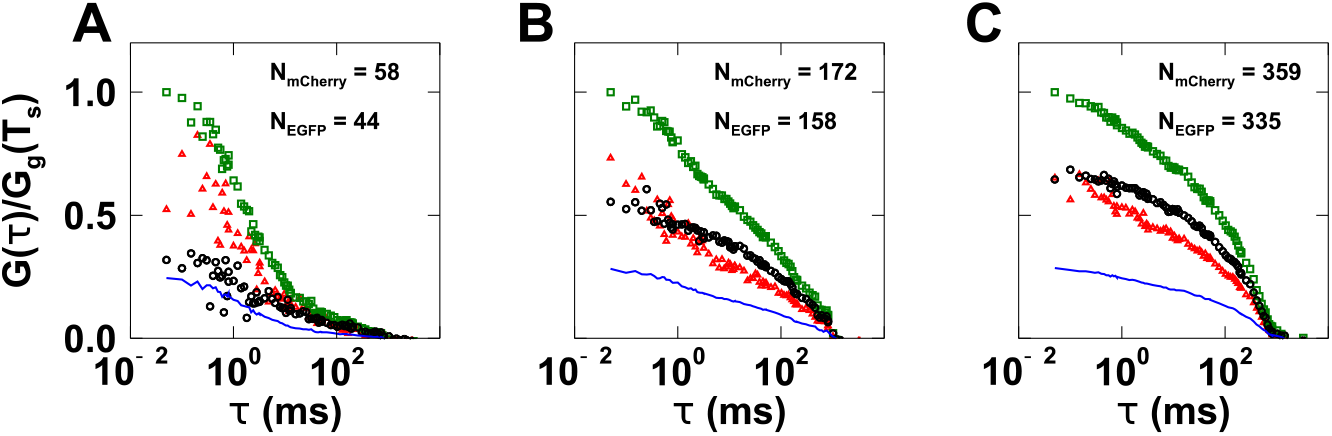
ACF and CCF analysis of DC FFS data collected in the NE of cells expressing SS-EGFP and SS-mCherry-KDEL at low (A), medium (B), and high (C) levels of protein expression. The number of proteins in the observation volume is given by *N*_EGFP_ and *N*_mCherry_. The red and green channel ACFs are shown with red triangles and green squares, respectively. The CCF is graphed as black circles and the calculated CCF from spectral crosstalk (CCF_ct_) is shown by the blue line. All correlation curves are divided by the amplitude of the ACF of the green channel, *G*_*gg*_(*T*_*S*_), with a lag time equal to the sampling time *T*_*S*_.

The unusual behavior of the experimental ACF of FFS data collected within the NE was traced to undulations of the nuclear membranes [14]. These membrane undulations introduce local volume fluctuations that modulate the fluorescence intensity signals received from SS-EGFP and SS-mCherry-KDEL (Fig. 1B). This additional fluctuation process not only affects the ACF, but also the CCF as the volume fluctuations lead to concomitant intensity variations in both the green and red channels. We previously demonstrated that the slow membrane undulations lead to biases in ACF analysis [14]. To overcome this challenge we replaced ACF with MSQ analysis and more recently with the improved tsMSQ algorithm [14,18]. Both methods have proven to be robust for measuring homotypic protein interactions within the NE of living cells [16,25]. To extend crosscorrelation analysis to DC FFS data obtained in the NE, we generalize single-color to dual-color tsMSQ. Two subscripts are added to distinguish the different tsMSQ functions. The SC tsMSQ from the green channel is identified by tsMSQ_gg_, while the crosscorrelation tsMSQ between the green and the red channel is described by tsMSQ_gr_ (Fig. 3). The autocorrelation tsMSQ_gg_ is calculated from the fluorescence intensity of a single channel using the standard tsMSQ algorithm (Fig. 3A). The crosscorrelation tsMSQ_gr_ follows the same procedure but utilizes the intensity traces of both channels (Fig. 3B).

**Figure 3.**
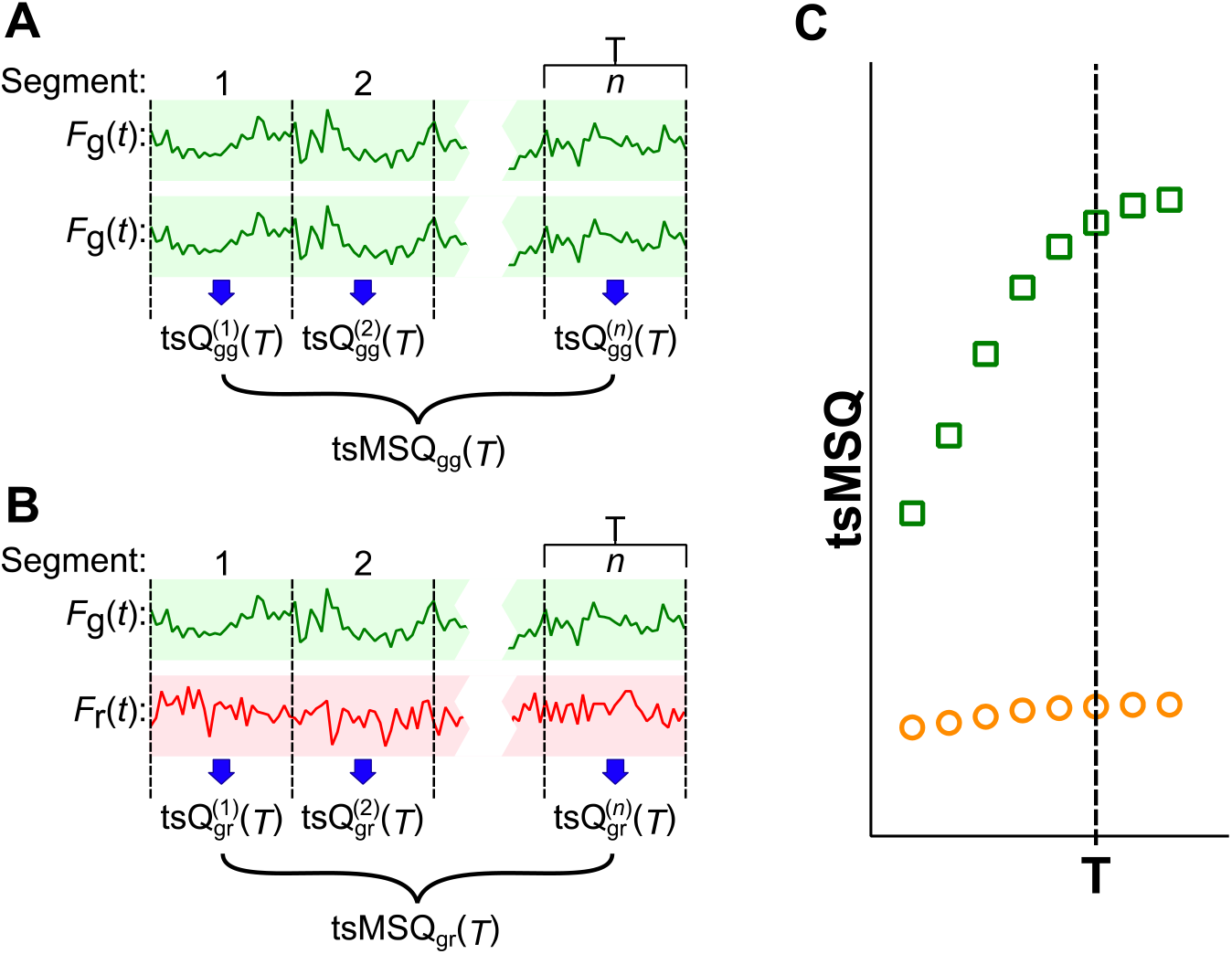
Conceptual illustration of the DC tsMSQ algorithm. A) The fluorescence intensity trace of the green channel, *F*_*g*_, is divided into *n* segments of period *T*. The time-shifted Q value of segment *i*, 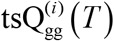, is calculated from *F*_*g*_ and a copy of *F*_*g*_ according to Eq. S3. Averaging over the time-shifted Q values determines the tsMSQ_gg_ for segment time *T*. B) The algorithm for calculating the crosscorrelation tsMSQ_gr_ follows the same procedure as described in panel A, but replaces the copy of *F*_*g*_ with the intensity of the red channel, *F*_*r*_. C) The autocorrelation tsMSQ_gg_ (green squares) and crosscorrelation tsMSQ_gr_ (orange circles) are constructed by repeating the procedures shown in panels A and B for a range of segment times *T*.

We reanalyzed the FFS data of SS-EGFP and SS-mCherry-KDEL taken in the NE (Fig. 2) using the experimental tsMSQ_gg_ (green squares) and tsMSQ_gr_ (red circles) curves (Figs. 4A and S4A-B). For two non-interacting proteins undergoing simple diffusion in the lumen, the crosscorrelation tsMSQ_gr_ should be entirely determined by the crosstalk *f*_ct_ from EGFP into the red channel and the membrane undulations of the NE, which contribute a correlation amplitude proportional to the average intensity [14]. Thus, the tsMSQ_gr_ curve for two non-interacting luminal proteins is predicted to be

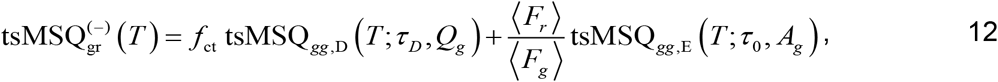

where tsMSQ_*gg*,D_(*T*;*τ*_*D*_, *Q*_*g*_) and tsMSQ_*gg*,E_(*T*;*τ*_0_, *A*_*g*_) are model functions (Eqs. 1 and 2) that describe the diffusion and exponential components of tsMSQ_gg,_, respectively, while 〈*F*_*i*_〉 represents the mean intensity in the *i*^th^ channel. The green-channel tsMSQ_gg_ was fit to a single species diffusion model with an exponential correlation term (Eq. 3) to account for nuclear membrane undulations, enabling the calculation of the predicted crosscorrelation 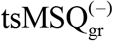 in the absence of interactions from Eq. 12 (blue line, Figs. 4A and S4A-B). The predicted 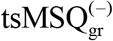 and the experimental tsMSQ_gr_ closely overlap, demonstrating the absence of protein interactions between luminal EGFP and mCherry-KDEL. These results demonstrate that tsMSQ_gg_ accurately predicts the cross-correlation 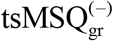 curve of a non-interacting pair of proteins within the NE.

**Figure 4.**
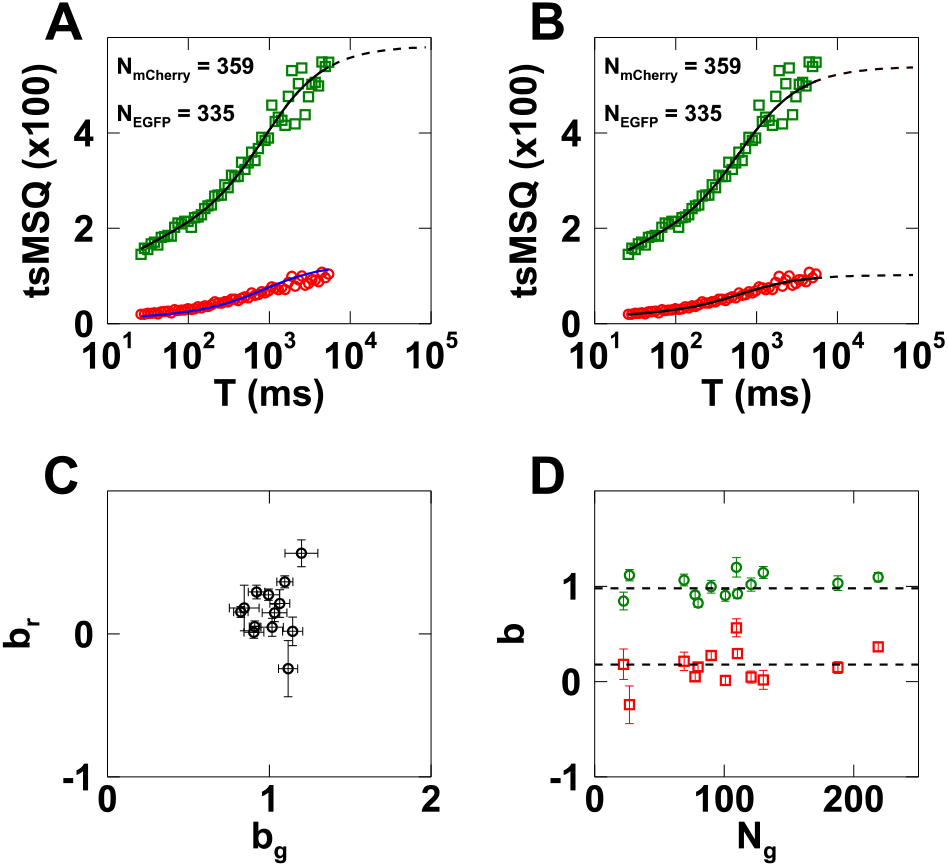
DC tsMSQ analysis of measurements performed in the NE of cells (*n* = 16) co-expressing SS-EGFP and SS-mCherry-KDEL. A) tsMSQ curves (symbols) with predicted 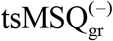 curve (blue line) derived from a fit to tsMSQ_gg_ (black line). The tsMSQ_gg_ and tsMSQ_gr_ curves are shown with green squares and red circles, respectively. B) tsMSQ curves (symbols) with fit to Eq. 7 (black). C) *b*_*r*_ vs. *b*_*g*_. D) *b*_*g*_ (green circles) and *b*_*r*_ (red squares) vs. *N*_*g*_ with means (dashed lines) and standard deviations of *b*_*g*_ = 1.0 ± 0.2 and *b*_*r*_ = 0.16 ± 0.13.

This initial validation of the theory prompted us to analyze FFS data with DC tsMSQ, which simultaneously describes the green-channel autocorrelation and the cross-correlation tsMSQ (Eq. 4), **tsMSQ** = [tsMSQ_gg_, tsMSQ_gr_]. We fit both tsMSQ curves to Eq. 7, which describes a single diffusion species in the presence of volume fluctuations. The fit and data are in very good agreement (Figs. 4B and S4C-D) with reduced chi-squared values close to one. The analysis was performed on *n* = 16 cells to collect fit parameters over a range of expression values. The amplitude **A** = (*A*_*g*_, *A*_*r*_) is expected to increase linearly with the fluorescence intensity, which agrees with the data (Fig. S5A). The characteristic time of the volume fluctuations is independent of concentration with a mean time of 0.3 s, closely mirroring results obtained in previous work (Fig. S6A) [14].

The fitted Q-vector 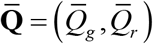 was converted into the normalized brightness **b** = (*b*_*g*_, *b*_*r*_) (Fig. 4C), as described in the Theory section. The values of the brightness plot scatter around **b** = (1,0), which is consistent for the SS-EGFP species containing one EGFP label and zero mCherry labels. These **b** values showed no dependence on the concentration of SS-EGFP (Fig. 4D). As discussed in the Theory section, DC tsMSQ was designed to identify HSP parameters. Thus, any purely red-emitting species is filtered out by HSP. As a consequence, we expected the non-interacting SS-mCherry-KDEL species to be invisible to the analysis, which is confirmed by the brightness plot. Consequently, the recovered diffusion times apply to the SS-EGFP species (Fig. S7A). Their values are concentration independent with a mean value of 1.7 ms, which is consistent with previous results [14,18].

As an additional control, we reanalyzed the cytoplasmic EGFP and mCherry data (Fig. S3) using DC tsMSQ with a fit to a single diffusing species (Eq. 5). The data and fit closely match (Fig. S8A-C), with the recovered HSP brightness values **b** = (*b*_*g*_, *b*_*r*_) centered near (1, 0) in a brightness plot, which is consistent with the EGFP species (Fig. S8D-E). The fitted diffusion time is concentration independent with a mean value of 0.8 ms (Fig. S7B), which agrees with previously published experiments [34].

After establishing that DC tsMSQ properly identifies non-interacting proteins within the NE and the cytoplasm, we next examined the tandem hetero-dimer SS-EGFP-SL-mCherry. This construct carries both EGFP and mCherry separated by a linker (SL) and mimics a strongly interacting pair of proteins forming a hetero-dimeric complex. As expected, DC FFS measurements of SS-EGFP-SL-mCherry in the NE produced a cross-correlation tsMSQ_gr_ curve that significantly exceeded the predicted baseline 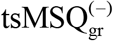 for non-interacting proteins (Fig. 5A). The DC tsMSQ curves were readily modeled by a fit to Eq. 7 (Fig. 5A) with reduced chi-squared values of 1.06. The HSP brightness values were centered around (1, 1), reflecting the presence of complexes with an average composition of one EGFP and one mCherry label as expected for the hetero-dimer (Fig. 5B). The diffusion time of the hetero-dimer identified by fitting is concentration independent (Fig. S7C). The **A** and *τ*_0_ values obtained for the volume fluctuations agree with the expected behavior (Figs. S5B, S6B). These results demonstrate that DC tsMSQ can be used to accurately identify the presence of hetero-dimeric protein complexes in the NE of living cells.

**Figure 5.**
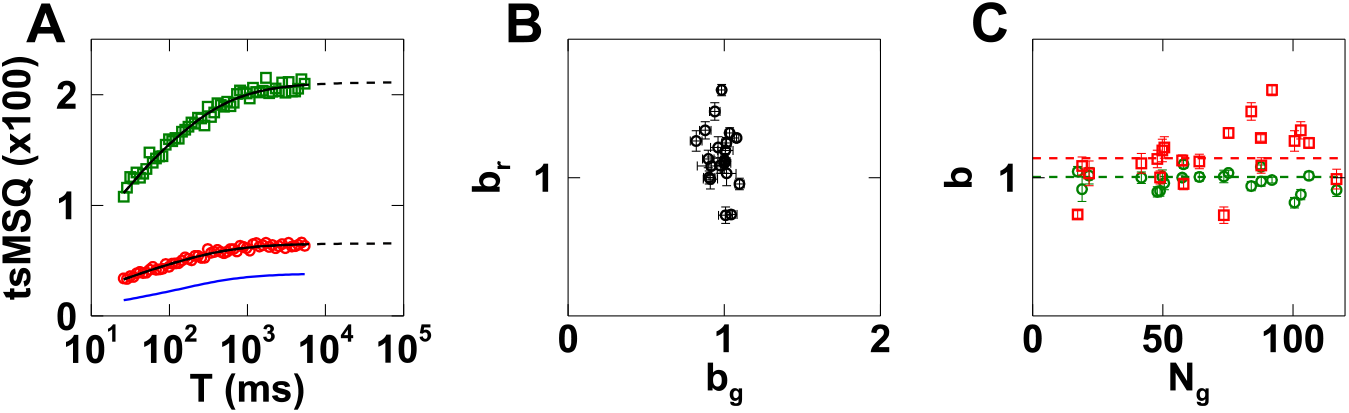
DC tsMSQ analysis of measurements obtained in the NE of cells (*n* = 24) expressing SS-EGFP-SL-mCherry. A) tsMSQ curves (symbols) with fits (black lines) and predicted 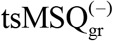 curve (blue line). The tsMSQ_gg_ and tsMSQ_gr_ curves are shown with green squares and red circles, respectively. B) *b*_*r*_ vs. *b*_*g*_. C) *b*_*g*_ (green circles) and *b*_*r*_ (red squares) vs. *N*_*g*_ with means (dashed lines) and standard deviations of *b*_*g*_ = 0.96 ± 0.06 and *b*_*r*_ = 1.1 ± 0.3.

While the fluorescence intensity of luminal proteins is affected by volume fluctuations in the NE (Fig. 1B), the fluorescence intensity of membrane-associated proteins is not affected by these volume fluctuations as demonstrated in previous work using SC FFS [14]. To test this difference between luminal and membrane-bound proteins in the context of DC tsMSQ, we performed DC FFS experiments in the NE of cells expressing the membrane-bound SS-EGFP-torsinA^NTD-2xLeu^ and the luminal SS-mCherry-KDEL. Since torsinA^NTD-2xLeu^ is a transmembrane domain [35], its presence ensures that EGFP is anchored to the nuclear membrane.

While the fluorescence signal generated by the luminal SS-mCherry-KDEL includes volume fluctuations, the fluorescence signal generated from the membrane-bound SS-EGFP-torsinA^NTD-2xLeu^ does not. Since DC tsMSQ filters out any purely red fluorescing species, the data are expected to only contain the membrane-bound green fluorescing species. Consequently, we fit the DC tsMSQ curves to a model containing only a diffusing species and no exponential process (Eq. 5), which agreed well with the data 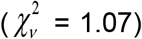. The fit recovered HSP brightness values **b** that are clustered around (1, 0) (Fig. 6B) and show no concentration dependence (Fig. 6C), which is consistent with SS-EGFP-torsinA^NTD-2xLeu^ being monomeric. The diffusion times for SS-EGFP-torsinA^NTD-2xLeu^ (Fig. S7D) have a mean and standard deviation of 16 ± 4 ms, which is in agreement with previously published results for this construct [14].

**Figure 6.**
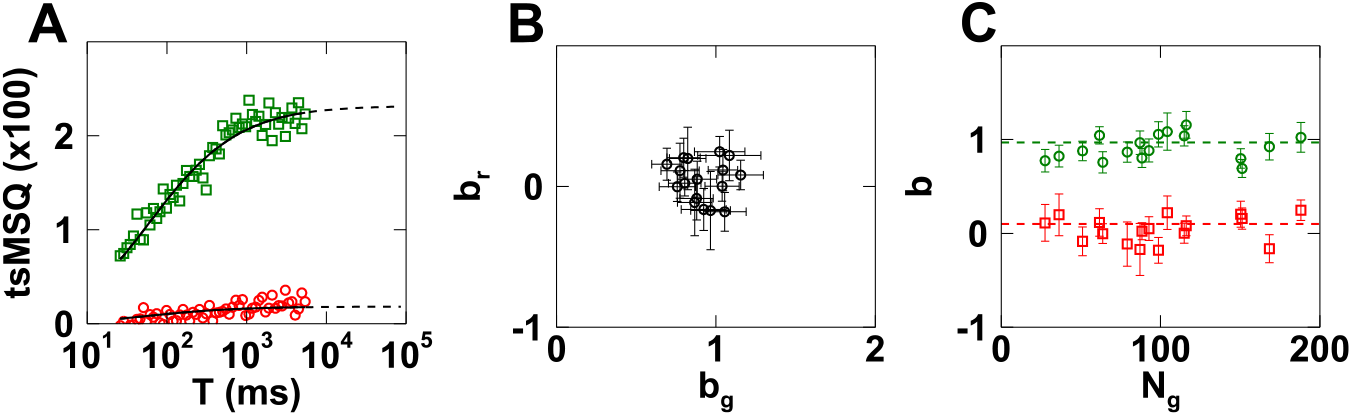
DC tsMSQ analysis of measurements performed in the NE of cells (*n* = 20) co-expressing SS-EGFP-torsinA^NTD-2xLeu^ and SS-mCherry-KDEL. A) tsMSQ curves (symbols) with fits (black lines). The tsMSQ_gg_ and tsMSQ_gr_ curves are shown with green squares and red circles, respectively. B) *b*_*r*_ vs. *b*_*g*_. C) *b*_*g*_ (green circles) and *b*_*r*_ (red squares) vs. *N*_*g*_ with means (dashed lines) and standard deviations of *b*_*g*_ = 0.97 ± 0.18 and *b*_*r*_ = 0.08 ± 0.12.

We next used DC tsMSQ to study the assembly of LINC complexes composed of SUN2 and nesprin-2 *in vivo* by investigating the ability of the SUN2 luminal domain to form a heterotypic interaction with the luminal domain of nesprin-2. Based on previously published *in vitro* biochemical and structural studies, it is expected that a homo-trimer of SUN2 interact with three nesprin-2 KASH peptides to form a hetero-hexamer [21,22]. Moreover, SUN2 homo-trimerization was shown to be critical for KASH-binding [21,36]. To begin to test this model of LINC complex assembly in the NE of living cells we first performed measurements on cells co-expressing mCherry-SR-KASH2 with SS-EGFP-SUN2^595-731^. The SUN domain (SUN2^595-731^) contains the KASH binding sites and has previously been shown to remain monomeric using SC tsMSQ [16]. Thus, the SUN domain permits us to directly test for the presence of monomer-monomer interactions with nesprin-2.

Fitting the DC tsMSQ curves to Eq. 7 agreed well with the data 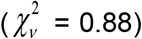 (Fig. 7A). A plot of *b*_*r*_ vs. *b*_*g*_ revealed that the data were clustered around (1, 0) indicating a non-interacting monomeric EGFP-labeled protein species (Fig. 7B). Neither *b*_*g*_ nor *b*_*r*_ showed any concentration dependence and both have means consistent with SS-EGFP-SUN2^595-731^ being unable to interact with mCherry-SR-KASH2 (Fig. 7C). In addition, the values of *τ*_*D*_, **A**, and *τ*_0_ all agreed with the expectation of a non-interacting SS-EGFP-SUN2^595-731^ monomer (Figs. S7E, S5C, S6C). Taken together, these results support the model in which the luminal domain of nesprin-2 is unable to interact with monomers of the SUN2 luminal domain.

**Figure 7.**
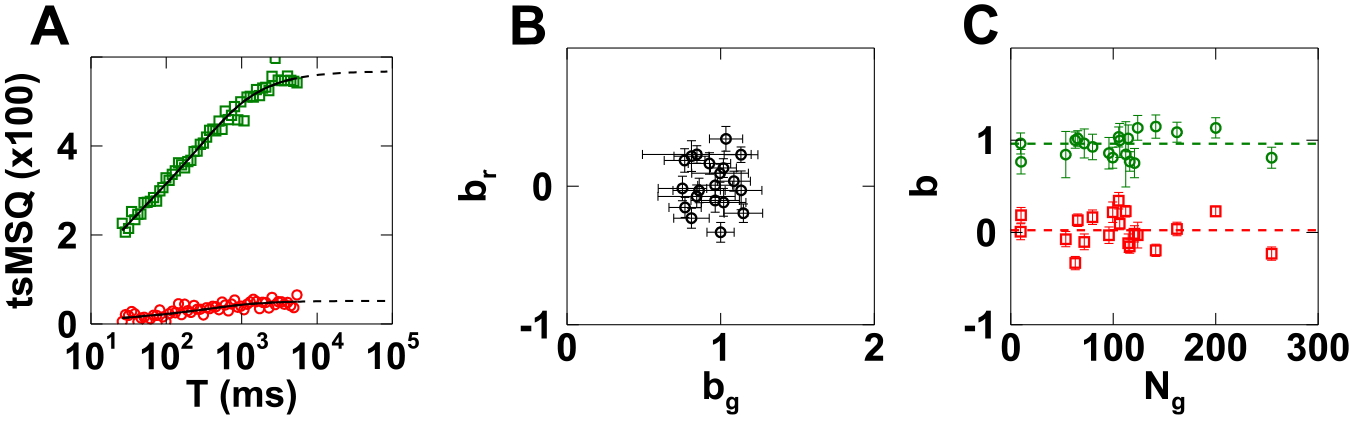
DC tsMSQ analysis of measurements performed in the NE of cells (*n* = 20) co-expressing SS-EGFP-SUN2^595-731^ and mCherry-SR-KASH2. A) tsMSQ curves (symbols) with fits (black lines). The tsMSQ_gg_ and tsMSQ_gr_ curves are shown with green squares and red circles, respectively. B) *b*_*r*_ vs. *b*_*g*_. C) *b*_*g*_ (green circles) and *b*_*r*_ (red squares) vs. *N*_*g*_ with means (dashed lines) and standard deviations of *b*_*g*_ = 0.96 ± 0.13 and *b*_*r*_ = 0.0 ± 0.2.

Unlike SS-EGFP-SUN2^595-731^, the SUN2 luminal domain-encoding SS-EGFP-SUN2^261-731^ homo-trimerizes in the NE of living cells as determined by SC FFS [18]. Thus, we expected that measurements performed in the NE of cells co-expressing SS-EGFP-SUN2^261-731^ and mCherry-SR-KASH2 would identify heterotypic interactions between these constructs. Proper analysis by DC tsMSQ of data obtained in cells expressing both proteins required the inclusion of two diffusing species (Eq. 8) to describe the data (Fig. 8A). The two diffusion times have means of 1.7 and 180 ms (Fig. S7F) which are consistent with a previous SC FFS study of SS-EGFP-SUN2^261-731^ [18].The tsMSQ_gr_ amplitude is significantly higher than the non-interacting prediction (Fig. 8A), indicating the presence of hetero-protein association. Plotting *b*_*r*_ vs. *b*_*g*_ from *n* = 53 cells revealed that both *b*_*g*_ and *b*_*r*_ increase together (Fig. 8B), implying that an increase in the average oligomeric state of SS-EGFP-SUN2^261-731^ is associated with an increase in the number of bound mCherry-SR-KASH2. This result supports the model that nesprin-2 interacts with homo-trimers of SUN2 within the NE. We further notice that *b*_*r*_ ≤ *b*_*g*_, indicating that at most one mCherry-SR-KASH2 can associate with each SS-EGFP-SUN2^261-731^ protein within the homo-trimer.

**Figure 8.**
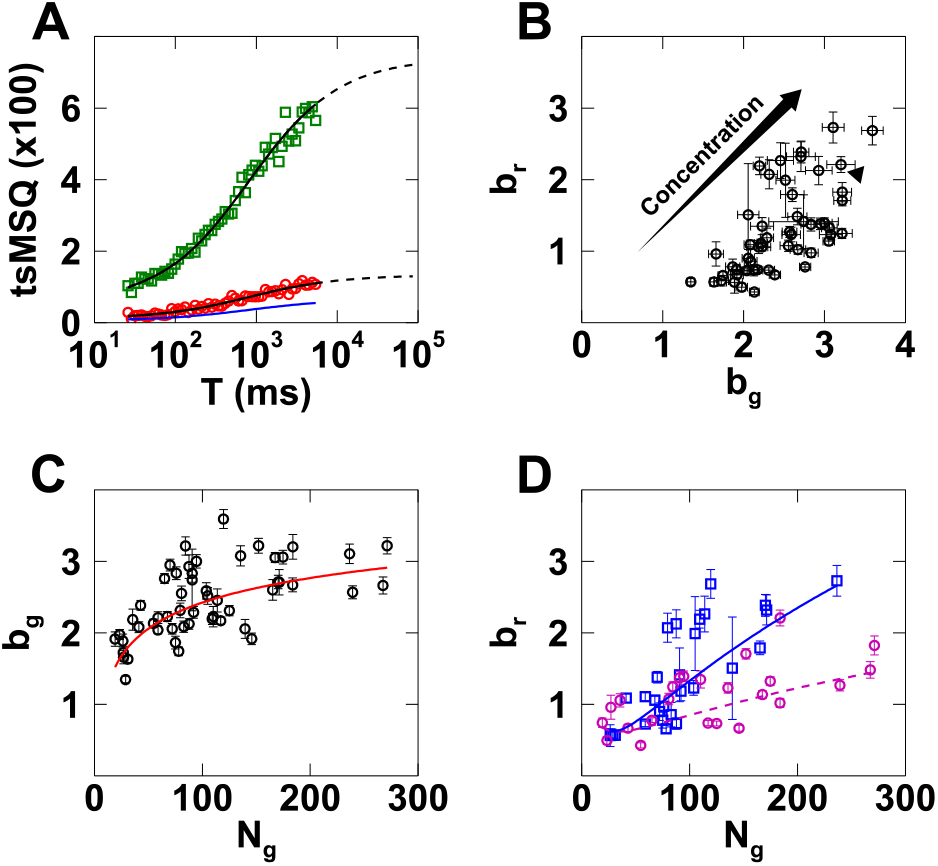
DC tsMSQ analysis of measurements performed in the NE of cells (*n* = 53) co-expressing SS-EGFP-SUN2^261-731^ and mCherry-SR-KASH2. A) tsMSQ curves (symbols) with fits (black lines) with 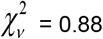 and predicted 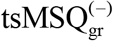 curve (blue line). The tsMSQ_gg_ and tsMSQ_gr_ curves are shown with green squares and red circles, respectively. B) *b*_*r*_ vs. *b*_*g*_ are correlated and increase with concentration as indicated. C) *b_g_* vs. *N*_*g*_ with a line provided to guide the eye. D) *b*_*r*_ vs. *N*_*g*_ separated by *N*_*r*_:*N*_*g*_ ratios greater (blue squares) or less (magenta circles) than 4:1, with lines provided to guide the eye.

These data show that *b*_*g*_ ranges between 1 and 3, signifying a limiting homo-trimeric state for the SS-EGFP-SUN2^261-731^ in the NE. The concentration dependence of the oligomerization of SS-EGFP-SUN2^261-731^ is visualized by a plot of *b*_*g*_ vs. *N*_*g*_ (Fig. 8C). The brightness increases with *N*_*g*_, approaching a homo-trimeric brightness state, which is identical to the behavior observed for SS-EGFP-SUN2^261-731^ in the absence of mCherry-SR-KASH2 [16]. We expect to observe an increase in *b*_*r*_ with *N*_*g*_, because the luminal domain of nesprin-2 is predicted to bind to homo-trimers of the SUN2 luminal domain, which are populated at higher *N_g_* (Fig. 8C). However, the amount of bound mCherry-SR-KASH2 also depends on its concentration, which varies significantly from cell to cell. Thus, the variability in the expression ratio of mCherry-SR-KASH2 to SS-EGFP-SUN2^261-731^ is responsible for the large scatter in the observed *b*_*r*_ values (Fig. 8D). To visualize the dependence of *b_r_* on the expression ratio of these two constructs, the data were separated into sets with an *N_r_*:*N_g_* above and below 4:1 (Fig. 8D). Data with a *N*_*r*_:*N*_*g*_ ratio above 4:1 exhibit a strong increase in *b*_*r*_ up to values approaching 3, while the lower ratio data only show a modest increase in *b*_*r*_ with *N*_*g*_. This reflects the relative reduction in mCherry-SR-KASH2 concentration between both data sets. Notably, the highest brightness values measured for *b*_*g*_ and *b*_*r*_ approach 3 (Figs. 3B-D), which suggests the formation of a SUN2-nesprin-2 hetero-hexamer *in vivo* as predicted by *in vitro* models [21,22].

## Conclusions

While CCF analysis has been widely used to identify heterotypic protein interactions from FFS measurements performed in cells [37], the presence of nuclear membrane undulations is a significant barrier for its application to FFS measurements performed in the NE. The local volume changes caused by nuclear membrane undulations lead to coupled changes in the emission intensity of both the red and green fluorescent labels (Fig. 1B), which result in a spurious crosscorrelation in the CCF, even when accounting for spectral crosstalk. The DC tsMSQ theory developed in this paper provides a comprehensive method that incorporates both spectral crosstalk and volume fluctuations, thus overcoming the issues present in CCF analysis of FFS data measured in the NE. We experimentally verified DC tsMSQ using model systems representing non-interacting protein pairs as well as hetero-dimeric protein complexes and recovered diffusion times and HSP brightness values. The label stoichiometry determined by HSP brightness matched the expected values of the control samples and reliably distinguished interacting from non-interacting proteins. Furthermore, the parameters of the volume fluctuation process agreed with our previously published SC tsMSQ results [14], thus providing additional support for the DC tsMSQ model.

Next, we used DC tsMSQ to investigate the assembly mechanism of LINC complexes, which are formed by the heterotypic interaction of the luminal domains of SUN and KASH proteins [38]. Previously published *in vitro* biochemical and structural studies revealed that a homo-trimer of the SUN2 luminal domain interacts with three nesprin-2 luminal domains [21,22]. Deep KASH peptide-binding grooves are formed at the interface of two adjacent SUN domains upon SUN2 luminal domain homo-trimerization; therefore, SUN2 homo-trimerization is believed to be a necessary precursor to the assembly of a SUN2-nesprin-2 hetero-hexamer [21,22].

To test this model *in vivo*, we looked for interactions between the luminal domain of nesprin-2 (mCherry-SR-KASH2) and two SS-EGFP-tagged SUN2 luminal domain constructs. We previously showed that the EGFP-tagged luminal domain of SUN2 (SS-EGFP-SUN2^261-731^) forms homo-trimers in the NE, while the SUN domain of SUN2 (SS-EGFP-SUN2^595-731^) remains monomeric [16]. As expected, we were unable to detect a heterotypic interaction between SS-EGFP-SUN2^595-731^ and mCherry-SR-KASH2 using DC tsMSQ. However, DC tsMSQ did detect a concentration-dependent heterotypic interaction between SS-EGFP-SUN2^261-731^ and mCherry-SR-KASH2 in the NE. The limiting HSP brightness values measured for these constructs suggest the formation of a SUN2-nesprin-2 hetero-hexamer *in vivo*, which agrees with the *in vitro* results described above [39].

These initial results provide a promising starting point for future quantitative studies of the mechanisms underlying the *in vivo* assembly of functional LINC complexes and their regulation. Combining quantitative modeling of the DC tsMSQ brightness data with targeted mutations that perturb the known SUN2-nesprin-1/2 binding sites should provide a comprehensive approach for investigating the assembly of SUN2-containing LINC complexes. In addition, DC tsMSQ offers a tool to investigate the formation of LINC complexes composed of lesser studied SUN proteins, such as SUN1, which forms higher-order oligomers than SUN2 [16]. While this paper has focused on the application of DC tsMSQ for characterizing the heterotypic interactions of LINC complex proteins, the development of DC tsMSQ offers a promising and general platform for future studies of hetero-protein complex formation in the NE of living cells.

## Author Contributions

J.H., G.W.G.L., and J.D.M. were responsible for experimental design. J.H. performed experiments. K.H. developed analytical tools. J.H., K.H., J.K., S.R.K., I.A., and J.D.M. performed analysis. J.H., K.H., G.W.G.L., and J.D.M. wrote the manuscript.

## Acknowledgements

This work was supported by the National Institutes of Health GM064589 (J.D.M, J.H., K.H., J.K., S.R.K., and I.A.) and GM129374 (G.W.G.L., J.D.M., and K.H.). We thank Cosmo A. Saunders for his help generating the DNA constructs used in this study.

